# High-Quality *De Novo* Genome Assembly for the Galápagos Endemic Lava Gull Using Oxford Nanopore Technologies

**DOI:** 10.1101/2025.07.21.665996

**Authors:** Jessica A. Martin, James B. Henderson, Vera de Ferran, Gabriela Pozo, Alice Skehel, Athena Lam, John P. Dumbacher, Jaime A. Chaves

## Abstract

High-quality reference genomes permit deeper investigation into species’ evolution and provide insight into species management and conservation. Next-generation sequencing technologies, such as Oxford Nanopore Technologies, allow researchers to generate high-accuracy long-read genetic data in real-time from anywhere in the world, increasing accessibility to sequence data. The lava gull (*Leucophaeus fuliginosus*), an endemic bird species of the Galápagos archipelago, is the world’s rarest gull with an estimated population of 300 to 600 individuals. Little genetic research has been done on this species due to its solitary nature and small population size. Here we present a chromosome-level reference genome assembly of an adult female lava gull, generated using ultra-long reads from the Oxford Nanopore Ultra-Long DNA Sequencing Kit and a PromethION 2 Solo device. Initial sequencing generated 1.78 million reads, consisting of 29.6 gigabases (Gbp), with a mean Q-score of 17.8 at an average 22.5x coverage. Our final assembly has a total length of 1.31 Gbp, with 450 scaffolds, and a scaffold N50 of 85.1 Mbp and contig N50 of 42.8 Mbp. The generation of a high-quality whole genome for the lava gull is an important step for investigation into the species’ phylogeography and population genetics.

## Introduction

High-quality whole genomes permit deeper investigation of a species’ evolution through intraspecific and interspecific comparison (Ryder, 2005), including studies in phylogeography and population genetics, which are critical for the assessment, monitoring, and management of biodiversity (Brito & Edwards, 2009; Genome 10K Community of Scientists, 2009; Paez et al., 2022; Romanov et al., 2009). Publicly accessible reference genomes enable researchers to align newly collected sequences with a species-specific reference (Pozo et al., 2024), which is crucial for a wide range of biological studies key to the conservation of a species (Brandies et al., 2019). This includes genome-wide discovery of single-nucleotide polymorphisms that support investigations into historical demography and population structure (Vlček et al., 2025), examinations into genetic diversity, genomic erosion, inbreeding, and genetic load (Díez-del-Molino et al., 2018; Van Der Valk et al., 2019), as well as disease susceptibility, and other conservation-related genetic traits (Díez-del-Molino et al., 2018; Fuentes-Pardo & Ruzzante, 2017). As sequencing technologies improve in portability, such as with Oxford Nanopore Technologies (ONT), they enable real-time genomic data generation in remote, biodiverse regions without sequencing infrastructure (Watsa et al., 2020). This helps overcome challenges in areas where biological sample export is restricted, such as the Galápagos Islands.

The Galápagos Islands, located 1,000 kilometers off the coast of Ecuador, are renowned for their role in shaping Darwin’s theory of evolution. Like other oceanic islands, the archipelago hosts many endemic species (Kier et al., 2009) that evolved from a combination of isolation, unoccupied niches, and a small number of founders. The resulting biodiversity makes the Galápagos important for global conservation efforts (Lack, 1947; D. Snow & Grant, 1988). For example, of the 199 bird species recorded, 47 are endemic (CDF 2025), but due to their small geographic distribution and population sizes, these species are highly susceptible to human disturbances (Benitez-Capistros et al., 2014; Chaves, 2018; Vitousek et al., 1997; Wiedenfeld & Jiménez-Uzcátegui, 2008). Evolving in the absence of invasive predators, native species are primarily threatened by introduced rats and cats (Carrion et al., 2011; Cruz & Cruz, 1987; Steadman, 1986). In addition, the archipelago has experienced an exponential increase in tourism since the 1960s, which developed land for urban infrastructure and agriculture (Epler, 2007; Walsh & Mena, 2016).

One Galápagos endemic species, the lava gull (*Leucophaeus fuliginosus*), is of particular concern due to increasing habitat loss and disturbance by urban activity (Jiménez-Uzcátegui et al., 2019). First described by John Gould (Gould, 1841), the lava gull is the rarest gull species in the world, with an estimated population of 300 to 600 individuals (Brinkhuizen & Nilsson, 2020; Grant et al., 2014) (Figure 1a). Lava gulls are adept scavengers and predators attracted to urban centers by human food and waste (Wiedenfeld & Jiménez-Uzcátegui, 2008), but their proximity to these areas makes them especially sensitive to anthropogenic disturbances (Jiménez-Uzcátegui et al., 2019; B. K. Snow & Snow, 1969) and pollutants in the environment (Muñoz-Pérez et al., 2023). Despite the lava gull’s small population size and classification as Vulnerable by the IUCN (BirdLife International, 2018), research on this species remains limited. No genomes are currently available for this species, limiting the ability to conduct high-resolution genetic analyses in a taxon that urgently requires it due to its imperiled conservation status.

**Figure 1.**
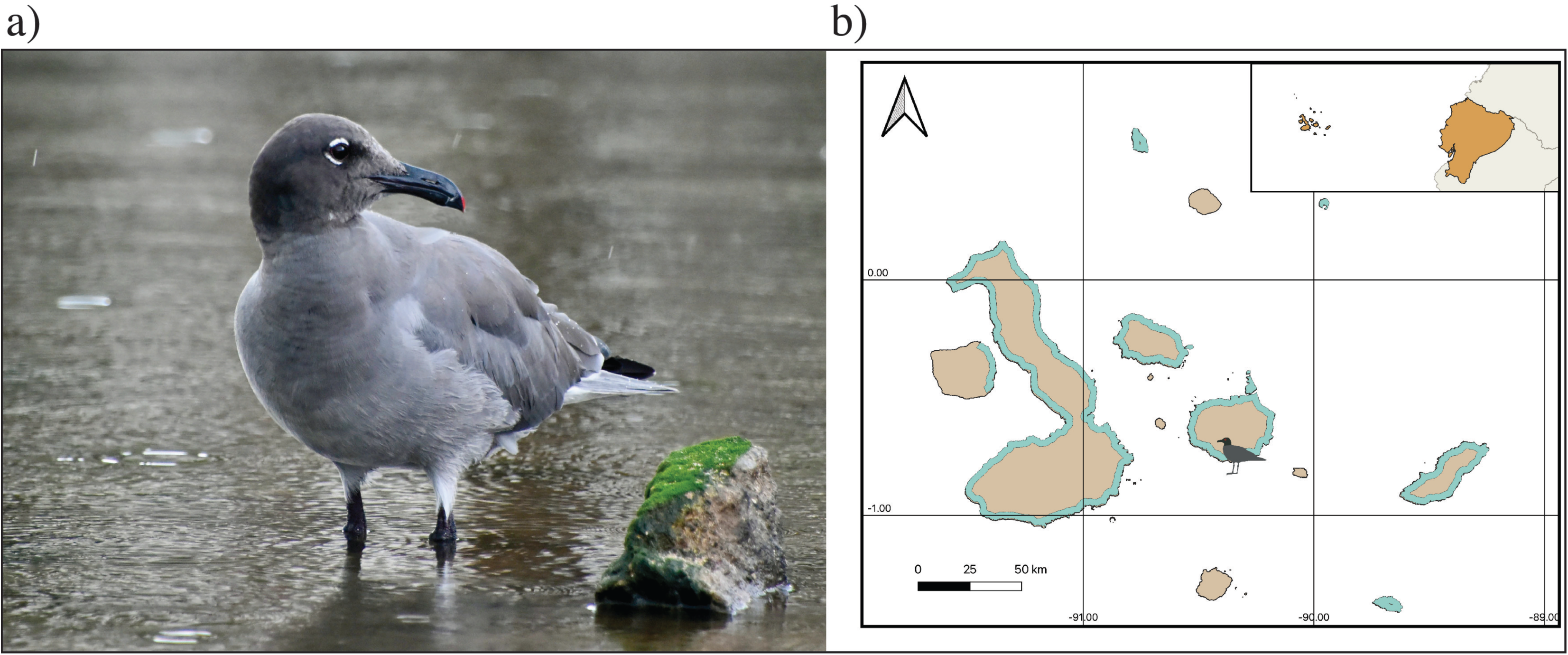
Lava Gull and Galápagos Islands. **a)** The lava gull is the rarest gull in the world and is endemic to the Galápagos archipelago. **b)** The Galápagos archipelago is located 1,000 kilometers off the coast of Ecuador (upper right inset). The sample used to create the lava gull reference genome was collected from an adult female gull at Puerto Ayora on Santa Cruz Island, as indicated by the lava gull icon on the map. Light blue on the Galápagos map indicates the lava gull’s current range (Brinkhuizen & Nilsson, 2020).

We present the first high-quality reference genome for the lava gull, a Galápagos-endemic and the world’s rarest gull species. All sequence data were generated in situ on the Galápagos Islands from a single female individual using Oxford Nanopore Technologies. We assess the accuracy and performance of two competing *de novo* assemblers, generate a complete mitogenome, and present an assembly with chromosomal resolution, including both avian sex chromosomes (W, Z). Finally, we compare the quality and completeness of our ONT genome (i.e., assembly metrics and synteny) to those of other gull (Laridae) genomes produced using different sequencing platforms.

## Methods and materials

### Sampling

A 0.4 ml blood sample was collected from the brachial vein using a 27-gauge needle and a 1ml syringe from an adult female lava gull on July 9, 2024, at the Muelle Municipal (Lat: 0°44’54.83”S, Long: 90°18’46.97”W) of Puerto Ayora, Santa Cruz Island, Galápagos (Figure 1b). The gull was caught using a baited remote-operated bowmen net and released once the blood sample and morphological measurements were taken. Upon collection, the sample was preserved in NAP buffer (Camacho-Sanchez et al., 2013) at the site and stored at −80°C upon return to the Galapagos Science Center (GSC) on San Cristóbal Island in the Galápagos.

### Sequencing Methods and Preparation

All lab work was performed on-site at the Galapagos Science Center, Puerto Baquerizo Moreno, San Cristóbal Island. DNA was extracted from nucleated blood using the Monarch HMW DNA Extraction Kit for Cells and Blood (New England Biolabs, Inc., Ipswich, MA), and its concentration was assessed with Qubit Fluorometric Quantitation (Thermo Fisher Scientific). A library was constructed using the Ultra-Long DNA Sequencing Kit SQK-ULK114 (Oxford Nanopore Technologies) and loaded on a PromethION 2 Solo (P2 Solo) device immediately after library completion. The P2 Solo was connected to a DELL Windows laptop with a 13th Gen Intel(R) Core(TM) i9-13950HX 2.20 GHz, 128 GB RAM, a 64-bit operating system, and x64-based processor for computational support and acquisition of the data onto the laptop drive. Sequencing was conducted using two PromethION Flow Cells FLO-PRO114M to generate POD5 format files that capture sequence signal data. Live basecalling requires GPU support and was not activated. Lab work and sequencing were performed following the manufacturer’s protocols with minor alterations outlined in the supplement (see Supplement: Sequencing Methods and Preparation). Subsequent analyses were performed at the California Academy of Sciences, San Francisco, CA, USA (CAS).

### Basecalling and Adapter Removal

For basecalling, POD5 files were copied to a desktop computer with GPU support. We then used the super-accurate model (SUP) in Dorado v0.7.4 (github.com/nanoporetech/dorado), the most accurate of the supplied models, and performed basecalling through minKNOW core v6.0.8 to generate 56 gigabytes of fastq files. Basecalling resulted in gzip fastq files with ∼4,000 reads each, which we combined into a single fastq file and eliminated reads less than 250 bp. This fastq file was used from the CAS high-performance computing (HPC) environment for all additional work. Basecalled reads were analyzed with NanoPlot v1.43.0 (De Coster et al., 2018) and a custom script (seq_summary_qscore_lens.sh, see Code Availability) to determine initial sequencing results, including number of reads, read length, read N50, and mean quality. Library adapters were removed using the program Porechop v0.2.4 (Wick et al., 2017) to generate trimmed fastq reads, which were assessed with the same programs.

### Genome Assembly

To generate initial information, including genome size, coverage, and heterozygosity levels, GeneScopeFK (github.com/thegenemyers/GENESCOPE.FK) was used with kmer size 21. GeneScopeFK is a command-line implementation of GenomeScope2 (Ranallo-Benavidez et al., 2020), with the same results, that creates a kmer histogram and GenomeScope2 plot figures.

Two genome assemblers were used to determine which produced the best *de novo* assembly. The first was generated with Flye v2.9.5 (Kolmogorov et al., 2019) using the `-scaffold` option and trimmed read input. The second was constructed with Hifiasm v0.24 (Cheng et al., 2021, 2025) using trimmed read input and the `--ont` option to enable its ONT read error correction. Assembly quality and completeness were assessed by standard metrics, including N50, L50, and the longest contig using in-house modified script asmstats.pl (see Code Availability), seqtk `telo` (github.com/lh3/seqtk), compleasm v0.2.6 (Huang & Li, 2023) with miniprot (Li, 2023), and BUSCO v5.4.7 (Simão et al., 2015) with metaEuk (Levy Karin et al., 2020) using the aves_odb10 lineage containing 8,338 orthologs. The two assemblies were compared using these results. Though both did well, their BUSCO scores led to the selection of the Flye result for contig-level assembly used for follow-on input.

To screen for contamination and leftover adapters, the Flye assembly was run through NCBI’s Foreign Contamination Screen (github.com/ncbi/fcs) programs FCS-adaptor and FCS-GX against taxonomy ID 328047. Mitochondrial contaminants were identified and removed by blasting against the mitochondrial assembly (see Mitochondrial Assembly) to eliminate any mitochondrial reads present in the nuclear genome assembly. A minimum 80% read length mapping was selected to keep nuclear-integrated mitochondrial sequences (NUMTs) from being excluded.

### Chromosomal-Level Scaffolding by Reference

The *de novo* assembly was scaffolded with references from two species of Laridae, the lesser black-backed gull (*Larus fuscus*, GCA_963932225.1) (Lopez Colom et al., 2024) and the yellow-legged gull (*L. michahellis*, GCA_964199755.1) (Ramos et al., 2025), to create a mapped reference using RagTag scaffold v2.28 (Alonge et al., 2022). These two reference genomes were the highest quality chromosomal-level gulls available on NCBI at the time of assembly. The lesser black-backed gull lacked a W chromosome, so the yellow-legged gull was used to scaffold both the Z and W chromosomes. The purge-dups v1.2.5 (Guan et al., 2020) pipeline was run with the scaffolded assembly, identifying Haplotig, Repeat, and Junk elements in the assembly and removing them to create a new assembly fasta. The purged reference provided contig groupings, orientations, locations, and confidence intervals. Following this, gap filling between scaffolded contigs was attempted with quarTeT GapFiller v1.2.1 (Lin et al., 2023) using the trimmed reads input for the Flye assembly. Records shorter than 5,000 bases were removed and the assembly named bLeucoFulig_1.0.

### Repeat Analysis

Repeat models were identified by RepeatModeler v2.0.1 (Flynn et al., 2020) using RepeatScout (github.com/Dfam-consortium/RepeatScout), RECON (www.repeatmasker.org), LtrHarvest (Ellinghaus et al., 2008), and Ltr_retriever (github.com/oushujun/LTR_retriever) to identify transposable elements, DNA sequences that move locations within a genome, and other repeat motifs. Identified models were combined with curated Aves models from Dfam 3.8 (dfam.org/releases/Dfam_3.8). The combined models and the assembly file were inputs to Repeatmasker v4.1.5 (Smit et al. 2015), which outputs a table categorizing repeats and their lengths. Repeatmasker also outputs a soft-masked repeat version of the assembly used for subsequent analysis. This soft-masked version identifies the repeats with lowercase letters as opposed to a hard-masked version, which replaces repeats with Ns. Identification and masking of repeats prior to conducting gene annotation precludes their inclusion in analyzed sequences.

### Genome Annotation

The soft-masked genome was structurally annotated using *ab initio* gene modeling program BRAKER3 v3.0.8 (Gabriel et al., 2024) in EP mode since we had no specimen RNA-seq data. This uses cross-species protein homology to determine intron and exon structure and predicts genes with GeneMark-EP+ and AUGUSTUS (Brůna et al., 2021). We used OrthoDB v11 vertebrate proteins for the homology set (Kuznetsov et al., 2023). BRAKER3 was run with `--busco_lineage=aves` added to invoke TSEBRA (Gabrielli et al., 2024), which uses compleasm to enhance recovery of BUSCO proteins. This resulted in gff3 annotation, coding sequence DNA, and protein sequence amino acid files. From these BRAKER3 output files, gene models lacking a start and a stop codon were removed, as were gene models nested within another gene model.

Functional annotation began with InterProScan v5.72-103.0 (Jones et al., 2014), identifying protein domains within the amino acid sequences. Then DNA and amino acid sequences were queried in several GenBank databases, using blastn v2.15.0 with the nt database (built Dec 22, 2023), blastp v2.15.0 with the uniprot_sprot database (built Dec 31, 2023), and diamond blastp v2.1.10 (Buchfink et al., 2021) with TrEMBL and nr databases (both built Dec 20, 2023). Additionally, we searched the eggNOG v5.0.2 database through eggNOG-mapper v2.1.11 (Cantalapiedra et al., 2021). InterProScan protein domains and gene ontology were included in the gff3 annotation and sequences files for each gene model, as was the functional annotation description from the lowest eValue, maximum 1e-10, for each gene from blast or eggNOG results. BUSCO was used to evaluate the resulting retained gene models with the aves_odb10 lineage in its protein mode. The gene models were further evaluated with OMArk v2.0.3 (Nevers et al., 2025) and the associated OMA LUCA.h5 database. OMArk employs Hierarchical Orthologous Groups (HOGs) to evaluate proteome completeness.

### Genome Comparison

The lava gull genome assembly was compared with three other gull species currently available on GenBank: lesser black-backed gull, yellow-legged gull, and laughing gull (*Leucophaeus atricilla*, GCA_045784845.1) (Petersen 2024). The comparison of the completeness of these assemblies was assessed using the date they were published to GenBank, the assembly level, the synteny, sex chromosomes availability, their size, the number of complete and missing BUSCOs, and assembly metrics including: number of contigs and scaffolds, longest contig and scaffolds, contig and scaffold N50 and L50, contig and scaffold N90 and L90, auN50, and synteny visualized with Circos plots generated from two of the genomes’ fasta files and BUSCO results.

### Mitochondrial Assembly

Mitochondrial sequences were pulled from the trimmed reads used with Flye and assembled using HiFiMiTie v0.08 (github.com/calacademy-research/HiFiMiTie), a long-read mitochondrial assembler. Assembly of the mitochondrial genome was performed concurrently with the Flye nuclear assembly to allow for the identification and removal of potential mitochondrial contaminants from the nuclear genome. Briefly, sequencing reads were compared against avian entries in the NCBI mitochondrial database using BLAST. Reads with ≥ 50% sequence coverage were retained for mitochondrial genome assembly, annotation, and subsequent analysis of heteroplasmy and tandem repeat structure within the control region. The Aves taxonomic identification was used to determine a canonical tRNA starting point for the sequence and the mitochondrial genetic code. MITOS2 (Bernt et al., 2013; Donath et al., 2019) was used to create a consensus annotation to confirm the annotation produced in HiFiMiTie. A visual representation of the circular genome was generated in Geneious Prime v2025.1.3 (www.geneious.com) using the mitochondrial fasta file and the annotation gff file.

## Results and Discussion

### Sequencing

Sequencing generated 1.78 million basecalled reads comprising 29.6 Gbp. Reads had a mean length of 16.6 kbp and 17.8 mean quality score. This sequencing output represents 22.5x average coverage for the species based on an estimated genome size of 1.31 Gbp (Table S2). This 22.5x coverage obtained from the lava gull basecalled ONT reads is lower than other avian genome assemblies sequenced using differing sequencing platforms, generated coverages of 36.07x, 37x, 79x, and 117x coverage (Table S2). Though the coverage of our resulting lava gull assembly had two-thirds the coverage of the lowest other assembly, the quality and completeness were comparable to other assemblies sequenced at a higher coverage.

### Genome Assembly

The Flye assembly from 29.4 Gbp of trimmed reads has a total size of 1.35 Gbp, consisting of 1,363 contigs and 1300 scaffolds. The assembly has a contig N50 of 37.1 Mbp and a contig L50 of 12. The longest contig was 100 Mbp, and the longest scaffold was 141 Mbp. The HiFiasm assembly of trimmed reads has a total size of 1.30 Gbp, comprising fewer contigs (n=671) (Table 1). The assembly had a contig N50 of 41.2 Mbp and a contig L50 of 9 (Table 1). The choice of the Flye assembly over Hifiasm for the lava gull contig-level assembly is contrary to the results from the assembly of the Galápagos petrel (*Pterodroma phaeopygia*), which was sequenced using identical laboratory and sequencing protocols. However, with the Galápagos petrel data, Hifiasm performed better or the same in each of the sequencing metrics; in particular, both assemblies had only one missing BUSCO (Sessi et al., 2025). The difference in Hifiasm result between the two taxa may lie in the Galápagos petrel sequencing, at 36x, having 14x increased coverage over the lava gull input of 22x. This was due to the lava gull single sequencing library, which generated 1.78 million reads, as opposed to the Galápagos petrel sequencing, which used two libraries and generated 4.10 million reads (Table S2).

**Table 1-.** Sequencing results from Flye v2.9.5 and Hifiasm v0.24 contig-level assemblies. The Flye assembly has just one missing BUSCO, while the Hifiasm assembly misses 15.

| Metric | Flye v2.9.5 | Hifiasm v0.24 |
| --- | --- | --- |
| Number of Scaffolds | 1,300 | 671 |
| Longest Scaffold | 141 Mbp | 120 Mbp |
| Scaffold N50 / L50 | 38.4 Mbp / 10 | 41.2 Mbp / 9 |
| Number of Contigs | 1,363 | 671 |
| Total Contig Length | 1.35 Gbp | 1.30 Gbp |
| Longest Contig | 100 Mbp | 120 Mbp |
| Contig N50 / L50 | 37.1 Mbp / 12 | 41.2 Mbp / 9 |
| Contig N90 / L90 | 1.34 Mbp / 74 | 1.40 Mbp / 73 |
| auN50 | 52.1 Mbp | 50.0 Mbp |
| BUSCO counts, n=8,338 | C: 8,334, D: 17, F: 3, M: 1 | C: 8,316, D: 17, F: 7, M: 15 |
| BUSCO percentages, n=8,338 | C: 99.95%, F: 0.04%, M: 0.01% | C: 99.74%, F: 0.08%, M: 0.18% |

The Flye assembly was scaffolded using both the lesser black-backed gull and the yellow-legged gull assemblies, generating a 1.35 Gbp genome made up of 1,187 scaffolds, with the longest scaffold increasing to 218 Mbp and scaffold N50 increasing from 38.4 Mbp to 84.8 Mbp. Purge_dups yielded an assembly of 1.31 Gbp with 628 contigs, removing 737 contigs, 532 of which were identified as either Highcov (n=41), Junk (n=72), or Repeat (n=419). Gap filling closed 68 gaps, adding 1.07 Mbp bases to our genome assembly. This left 84 remaining gaps, all of which had flanking regions too short to use. After gap filling, 26 contig records less than 5,000 bases were removed.

### Final Scaffolded Assembly

The final chromosomal-scale scaffolded assembly resulted in a genome size of 1.31 Gbp (Figure 2a) and was named bLeucoFulig_1.0. It consists of 450 scaffolds with a mean length of 2.92 Mbp. Scaffold N50 length is 85.1 Mbp with L50 count 5. The assembly is also made up of 533 contigs with a mean length of 2.46 Mbp. Contig N50 length is 42.8 Mbp at L50 count 9 (Table 2, column 2 lava gull). Comparing long contigs and scaffolds to the chromosome-level assembly of the lesser black-backed gull and yellow-legged gull, there were 34 pseudo-chromosomes in the lava gull genome. Of these 34 pseudo-chromosomes, nine have one telomere, including the Z chromosome, which was the fifth-longest chromosome in the genome with a length of 85.1 Mbp. This length is comparable to other avian Z chromosomes (Figure S4). We named autosomal sequences to match the numbers assigned to chromosomal records in the reference scaffolds. These results are similar to other publicly available chromosomal gull assemblies, ranging from 32 to 35 chromosomes. The use of reference-based scaffolding, as used here with the lesser black-backed gull and the yellow-legged gull, is an effective way of producing chromosomal-level scaffolds in birds (Jorquera et al., 2025; Shin et al., 2025) and mammals (Alvarez-Costes et al., 2025), assuming conservation of synteny (Takagi & Sasaki, 1974; Waters et al., 2021). From the 8,338 BUSCO genes, BUSCO identified 99.95% complete genes (single-copy: 99.75%, duplicated: 0.20%), 0.04% fragmented, and 0.01% missing. Of the 17 duplicated BUSCOs, 11 of these duplications occur as a single copy on each of the Z and W chromosomes (i.e., they are gametologs), indicating there is a high degree of paralogy between the two sex chromosomes and suggesting that they were appropriately assembled and counted. This is similarly reported in the yellow-legged gull, which has 22 duplicated BUSCOs, 14 of which occur on the Z and W chromosomes. With 99.95% complete BUSCO genes, this assembly contains a high number of conserved genes, indicating a highly complete assembly (Huang & Li, 2023).

**Figure 2.**
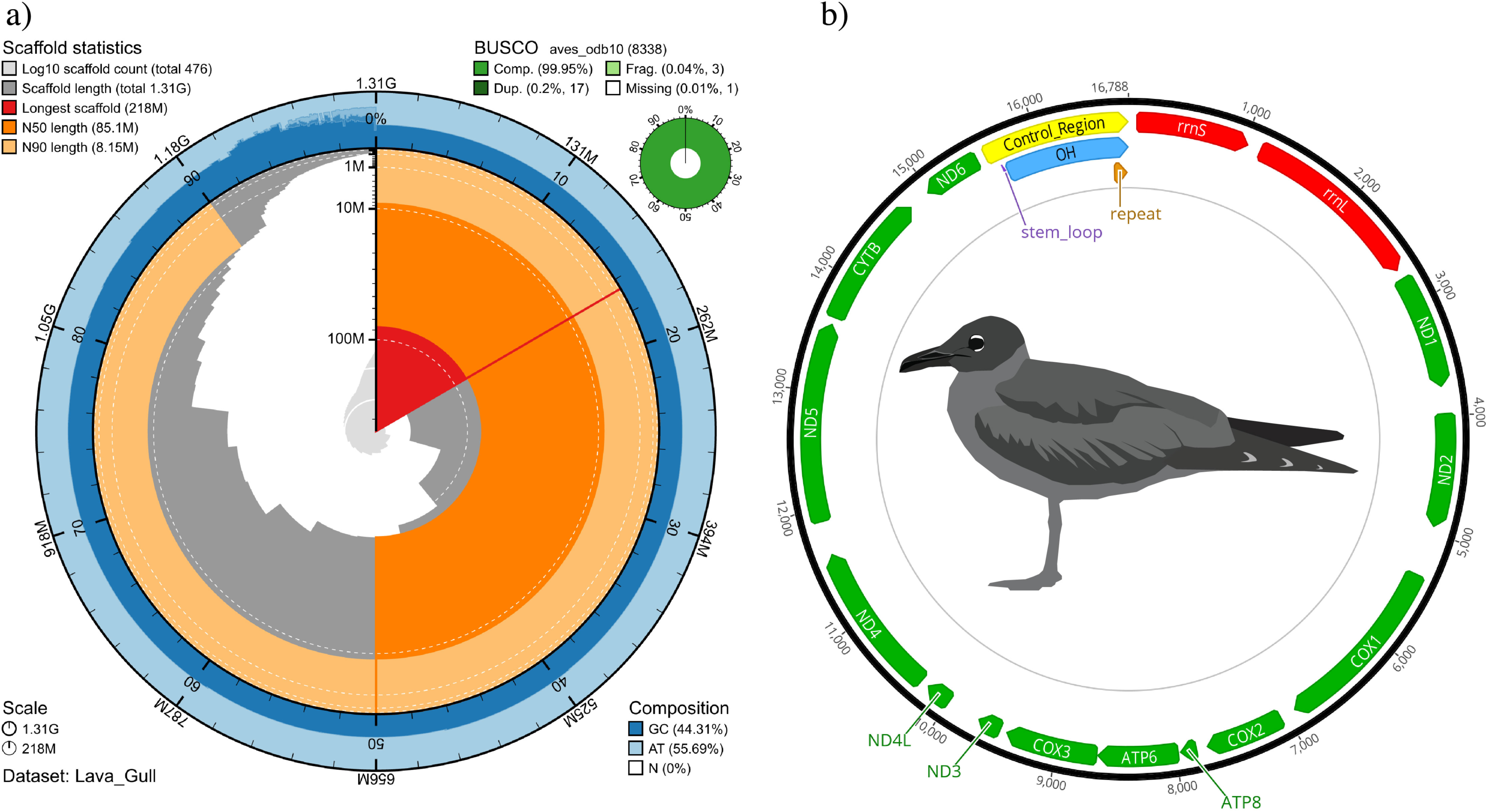
Snail and mitogenome plots for the Lava Gull. **a)** BlobToolKit snail plot depicting the assembly metrics for the lava gull genome. The circumference of the circle represents a genome size of 1.31 Gbp. The pale orange indicates scaffold N90 length of 8.15 Mbp, the deeper orange indicates scaffold N50 of 85.1 Mbp, red indicates the longest scaffold of 218 Mbp, and the light blue and dark blue on the perimeter indicate the AT and GC composition, respectively. The top right circle depicts the results from the BUSCO analysis, illustrating that the assembly was highly complete, containing 99.95% complete BUSCOs and only one missing and three fragmented BUSCOs. **b)** The mitochondrial genome assembly of the lava gull is made up of 16,788 bases and contains 13 protein-coding genes, illustrated in green, and two rRNAs, denoted in red.

**Table 2-.** Genome assembly comparison between the lava gull, laughing gull, lesser black-backed gull, and yellow-legged gull using assembly metrics including: number of contigs and scaffolds, longest contig and scaffold, contig and scaffold N50 and L50, which sex chromosomes are present and their lengths. Bolded values indicate the best value in each metric. BUSCO (Benchmarking Universal Single-Copy Orthologs) provides measures for quantitative assessment of genome assembly: C: complete; D: duplicated; F: fragmented; M: missing.

| Metric | lava gull<br>(*Leucophaeus fuliginosus*) | laughing gull<br>(*Leucophaeus atricilla*) | lesser black-backed gull<br>(*Larus fuscus*) | yellow-legged gull<br>(*Larus michahellis*) |
| --- | --- | --- | --- | --- |
| Date | This genome | Dec 6, 2024 | Feb 7, 2024 | Jul 14, 2024 |
| Genome Size | 1.31 Gbp | 1.18 Gbp | 1.32 Gbp | 1.41 Gbp |
| Genome Coverage | 22.5x | 160x | 30x | 68x |
| Number of Scaffolds | **450** | 125,582 | 1,367 | 618 |
| Longest Scaffold | 218 Mbp | 269 kbp | 218 Mbp | **224 Mbp** |
| Scaffold N50 / L50 | 85.1 Mbp / **5** | 40.4 kbp / 10,320 | 86.5 Mbp / **5** | **87.7 Mbp / 5** |
| Scaffold N90 / L90 | **8.15 Mbp / 21** | 4.99 kbp / 42,689 | 3.51 Mbp / 25 | 2.90 Mbp / 29 |
| Number of Contigs | **533** | 131,549 | 1,904 | 1,118 |
| Longest Contig | **141 Mbp** | 221 kbp | 17.2 Mbp | 17.7 Mbp |
| Contig N50 / L50 | **42.8 Mbp / 9** | 37.3 kbp / 10,756 | 3.57 Mbp / 106 | 4.22 Mbp / 98 |
| Contig N90 / L90 | **3.25 Mbp / 41** | 4.82 kbp / 45,749 | 421 kbp / 436 | 901 kbp / 369 |
| Contig auN50 | 101 Mbp | 39.4 kbp | **129 Mbp** | 99.8 Mbp |
| BUSCO n=8338 | **C:8334**, D:17, | C: 6828, D:7, | C: 8332, D: 7, | C: 8331, D: 22, |
|  | **F:3,M:1** | F:981, M: 529 | **F:5, M: 1** | F:4, M: 3 |
| BUSCO percentage | **C: 99.95%,**<br>**F:0.04%, M: 0.01%** | C: 81.89%,<br>F:11.77%, M: 6.34% | C: 99.93%,<br>F:0.06%, **M: 0.01%** | C: 99.92%,<br>F:0.05%, M: 0.04% |
| Assembly Level | Chromosome | Scaffold | Chromosome | Chromosome |
| Sex Chromosomes | Z, W (female) | N/A | Z (male) | Z, W (female) |
| Sex Chromosome<br>Sizes Z / W | 85.1 Mbp / 25.6Mbp | N/A | 214 Mbp / N/A | 87.7 Mbp / 26.9 Mbp |
| Sequencing<br>Platform | 22.5x ONT<br>Ultra-long | DIP-SEQ | 30x PacBio, Arima2<br>High-C data | 68x PacBio, Arima2<br>High-C data |

### Repeat Analysis and Annotation

RepeatMasker masked 209 Mbp or 15.89% of the 1.31 Gbp genome, comparable to other bird genomes available. Of the 209 Mbp of repeats, long interspersed nuclear elements (LINE) comprised 5.67%, long terminal repeat (LTR) elements 1.86%, and DNA transposons 0.56% (Figure S2, Table S12). Using kmer length 21, GeneScopeFK, a reimplementation of GenomeScope2, estimated a genome size of 1.15 Gbp, smaller than the assembled genome size of 1.31 Gbp. GeneScopeFK also identified 83.3 Mbp of repeats and estimated a genome heterozygosity of 0.38%. This underestimates the number of repeats by 125 Mbp, an underreporting commonly observed with this method; however, the estimated non-repeat length of 1.08 Gbp closely aligns with the 1.10 Gbp of non-repeats identified by RepeatMaster, suggesting general concordance between the two approaches for non-repeat content.

The BRAKER3 candidate gene model annotations built from the repeat-masked genome identified 24,768 genes and 26,902 mRNAs. Functional annotation and refinement yielded 21,808 gene models and 23,928 mRNAs (Table S14). From these 23,928 sequences, OMArk identified the input as belonging to Neognathae. Of the 11,061 conserved Hierarchical Orthologous Groups (HOGs) expected for this clade, 10,083 HOGs were found in these mRNAs. The proteome consisted of 21,808 proteins, of which 17,954 were consistent, 2,472 were inconsistent, and 1,382 were unknown (Figure S3, Table S15).

### Mitochondrial Genome

The mitochondrial genome was assembled from 37 adapter-trimmed reads using the HiFiMitie pipeline. The mitogenome contained 22 tRNAs, 2 rRNAs, and 13 protein-coding genes (Figure 2b), consistent with most vertebrate mitogenomes (Formenti et al., 2021). The assembled circular mitochondrial genome is 16,788 bp, similar in size to other gull species: 16,701 bp in the southern black-backed gull (*Larus dominicanus*) (Slack et al., 2007) and 16,788 bp in the black-tailed gull (*L*. *crassirostris*) (Kim & Park, 2016). The single control region consisted of 1,242 bp and contained a 7 bp consensus repeat motif repeated 16.6 times. Given that the Oxford Nanopore Technologies (ONT) sequencing platform is designed to target long reads during sequencing (store.nanoporetech.com/us/ultra-long-dna-sequencing-kit-v14.html) and with a sequencing mean read length of 16,586 bp in the present study, reads of mitochondrial origin are likely to be underrepresented or missed.

### Genome Comparison

The species used for the comparison include the laughing gull, which belongs to the same genus (*Leucophaeus*) as the lava gull, as well as the lesser black-backed gull and the yellow-legged gull, both in the genus *Larus.* The lesser black-backed gull and the yellow-legged gull were the most complete gull genomes available on GenBank, and the laughing gull was the only available genome in the same genus as the lava gull at the time of our lava gull assembly. Of the four, the lava gull genome assembly presented here has the fewest number of contigs at 533 and scaffolds at 450, the longest contig at 141 Mbp, the longest contig N50 at 42.8 Mbp, and the smallest contig L50 of 9 (Table 2).

The lava gull assembly also had the most complete BUSCOs at 8,334, and both the lava gull and the yellow-legged gull were missing only one BUSCO. This same BUSCO (28097at8782) was not present in this lava gull assembly, nor the three gull species used in this comparison, as well as two other species, the black-headed gull (*Chroicocephalus ridibundus*, GCA_963924245.1) (Colom et al., 2025) and the European herring gull (*Larus argentatus*, GCA_964417175.1) (www.genomeark.org/vgp-all/Larus_argentatus.html). This could indicate that this BUSCO is absent from the lineage; however, this requires further investigation into other Laridae genera. The large percentage of complete BUSCOs (99.95%) indicates that the lava gull assembly is highly complete, surpassing the quality of the other available *Leucophaeus* genome.

Though the initial sequence coverage of the lava gull, at 22.5x, was the lowest compared to other gull species (Table 2), we were able to generate a highly complete and contiguous reference genome, missing just one aves lineage BUSCO. This illustrates the utility and performance of the longer and more accurate ONT portable sequencing technology for genome assembly, even at a relatively low coverage and sequenced on-site in the Galápagos archipelago.

The contig synteny of the lava gull assembly was compared with the chromosomal-level assemblies for the lesser black-backed gull and the yellow-legged gull, which were both used for the reference-based scaffolding of the lava gull genome. The lava gull assembly had a high degree of synteny in comparison with both reference genomes (Figure 3). In the synteny assessment with the lesser black-backed gull, several BUSCOs located on the W chromosome of the lava gull were mapped to the Z chromosome of the lesser black-backed gull. This discrepancy resulted from the lack of a W chromosome from the male lesser black-backed gull; therefore, sequences from the lava gull’s W chromosome map to the Z chromosome in the absence of a corresponding W scaffold. The Z chromosome in the lesser black-backed gull, at length 214 Mbp and absence of BUSCO mapping in about 60% of the scaffold, is likely a misassembly. Also, an in-house script we use to complement seqtk telo, telomere_report.sh (see Code Availability), finds 26 telomere sequences mapping in this BUSCO desert. These ranged from 1,332 bp to 25,308 bp in length, starting at position 88.14 Mbp (Table S17).

**Figure 3.**
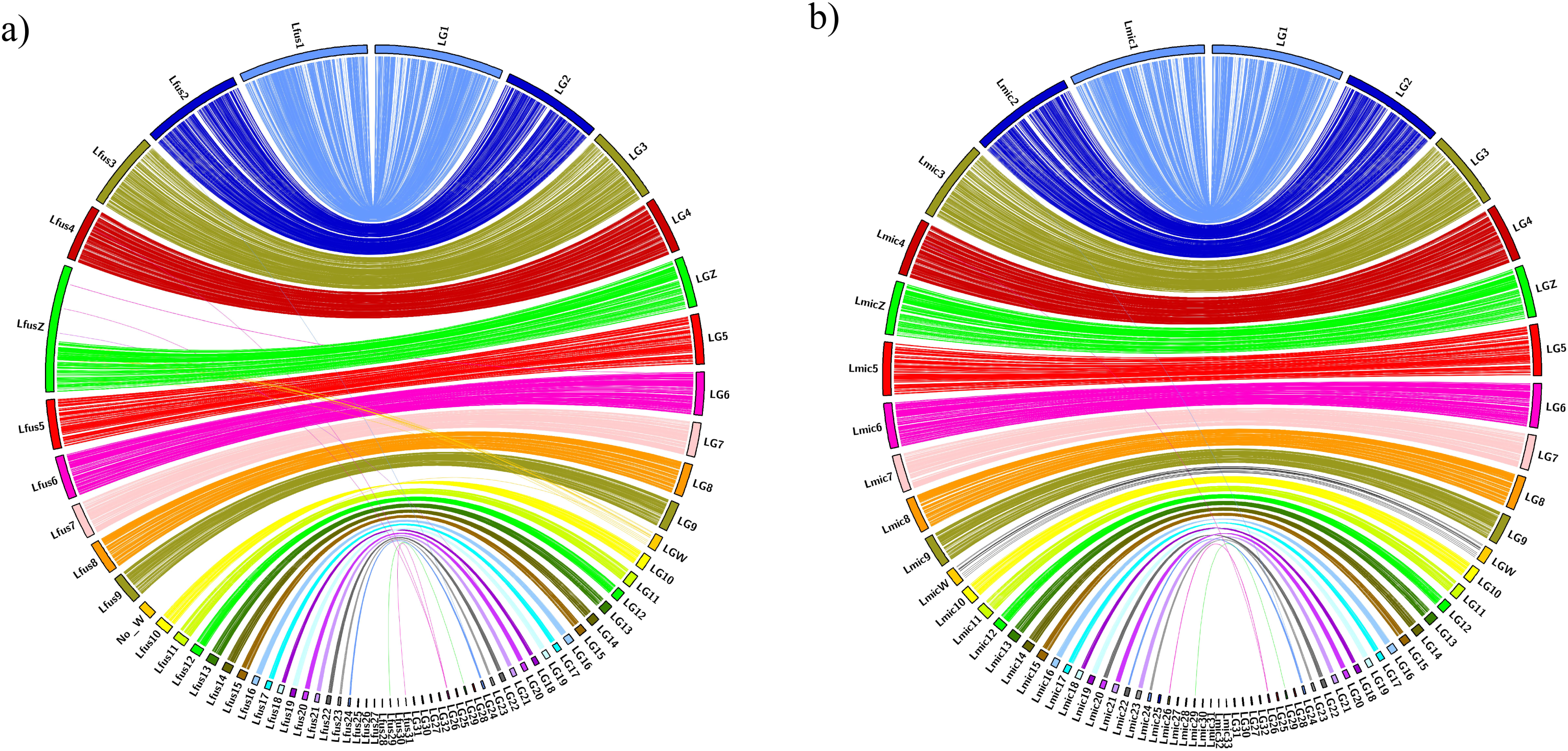
Synteny Between the Lava Gull Genome and the Lesser Black-Backed and Yellow-Legged Gulls. Circos plots illustrating synteny between the lava gull genome and **a)** the lesser black-backed gull and **b)** the yellow-legged gull. In each plot, the lava gull assembly is shown on the right and the comparative genome on the left.

## Conclusions

We present the first highly complete genome of a Galápagos endemic Charadriiformes species, assembled using Oxford Nanopore sequencing performed in the Galápagos Islands, and the first chromosomal-level assembly for the genus *Leucophaeus*. With a haploid genome of 1.31 Gbp, our assembly is similar to other gulls available on GenBank (range 1.2 Gbp to 1.4 Gbp), surpassing all previously published genomes for Laridae based on quality and completeness. Furthermore, our lava gull genome has the highest BUSCO completeness of 99.95% compared to other available Laridae genomes and makes available both sex chromosomes for this genus. This newly assembled genome will be immediately accessible to a wide range of biological studies that inform science-based conservation and management strategies. More broadly, our study demonstrates the feasibility of generating high-quality, chromosomal-level genome assemblies using portable sequencing technologies in remote field settings-opening new possibilities for real-time biodiversity genomics in regions where rapid, in situ data generation can have the greatest impact.

## Data Availability

All raw sequencing data are deposited at NCBI under PRJNA1275045, BioProject SAMN49010278, and BioSample SAMN49010278 : LVGU_060. The genome assembly is deposited under the same BioProject on NCBI and will be made public at the time of publication. It can be accessed using this link (reviewers’ only): https://dataview.ncbi.nlm.nih.gov/object/PRJNA1275045?reviewer=3bu7kcf60sgf0f7k39oj81j4u1

The publicly accessible datasets used in this study (lesser black-backed gull, yellow-legged gull, laughing gull, European herring gull, black-headed gull, little vermilion flycatcher, dark-eyed junco, and Galápagos petrel) can be found on GenBank and are cited within this paper.

## Code Availability

Custom scripts are in the github.com/jaimechaves76/GalapaGenomes-Lava-Gull-G3 GitHub repository. Scripts and programs starting from the adapter removal of the raw fastq files to the assembly annotation are located there, including seq_summary_qscore_lens.sh for read statistics, asmstats.pl for assembly statistics, basic_gff_stats.sh, telomere_report.sh, and script dependencies. The bioawk_cas program (github.com/calacademy-research/bioawk.CAS), an enhanced version of Heng Li’s bioawk (github.com/lh3/bioawk), is used by several scripts.

## Acknowledgments

We thank the Center for Comparative Genomics at the California Academy of Sciences for their assistance and training, the Galapagos Science Center for allowing us to conduct this work in their facility located on San Cristóbal-Galápagos, including Paúl Yepéz, Cristina Vintimilla, Gabriela Bautista, Jessenia Sotamba, and Sylvia Sotamba, and the Agencia de Regulación y Control de la Bioseguridad y Cuarentena para Galápagos (ABG), in particular, to Alberto Vélez. We would also like to thank the many field assistants who participated in fieldwork, including rangers from the Galápagos National Park, Andrea Loyola, veterinary students from NC-State University, and community members Andrea Villacres and Desire Sobreno. All samples were collected following approved protocols by the Directorate of the Galapagos National Park (permits PC 74-17 and PC-27-19). All material for the reference genome was accessed under MAE-DNB·CM-2016-0041 to JAC. All work was done under the Permiso de Acceso al Recurso Genético-Ecuador MAATE-DBI-CM-2022-0249. This genome is part of the *GalapaGenomes* project funded by NSF (BRC-BIO) to J.A.C. under San Francisco State University IACUC A2021-27. This work is part of J.A.M.’s master’s thesis, partially funded by the Student Enrichment Opportunities (SEO) at San Francisco State University (SFSU).

## Author Contribution

J.A.C. and J.P.D. designed the project; J.A.C. acquired funding for the project; A.L. provided critical training and guidance in the lab; J.A.M. and G.P. conducted the lab work and produced all the genetic data; J.B.H. and J.M. performed the bioinformatic analysis; J.A.M. wrote the manuscript with input from J.B.H., J.P.D., J.A.C., and V.F.; A.S. acquired funding for fieldwork and the biological sample. All authors checked and approved the final version of the manuscript.

